# Resting State Functional Connectivity Predicts Future Changes in Sedentary Behavior

**DOI:** 10.1101/2021.01.26.428161

**Authors:** Timothy P. Morris, Aaron Kucyi, Sheeba Arnold Anteraper, Maiya Rachel Geddes, Alfonso Nieto-Castañon, Agnieszka Burzynska, Neha Gothe, Jason Fanning, Elizabeth Salerno, Susan Whitfield-Gabrieli, Charles H. Hillman, Edward McAuley, Arthur F. Kramer

## Abstract

Information about a person’s available energy resources is integrated in daily behavioral choices that weigh motor costs against expected rewards. It has been posited that humans have an innate attraction towards effort minimization and that executive control is required to overcome this prepotent disposition. With sedentary behaviors increasing at the cost of millions of dollars spent in health care and productivity losses due to physical inactivity-related deaths, understanding the predictors of sedentary behaviors will improve future intervention development and precision medicine approaches. In 64 healthy older adults participating in a 6-month aerobic exercise intervention, we use neuroimaging (resting state functional connectivity), baseline measures of executive function and accelerometer measures of time spent sedentary to predict future changes in objectively measured time spent sedentary in daily life. Using cross-validation and bootstrap resampling, our results demonstrate that functional connectivity between 1) the anterior cingulate cortex and the supplementary motor area and 2) the right anterior insula and the left temporoparietal/temporooccipital junction, predict changes in time spent sedentary, whereas baseline cognitive, behavioral and demographic measures do not. Previous research has shown activation in and between the anterior cingulate and supplementary motor area as well as in the right anterior insula during effort avoidance and tasks that integrate motor costs and reward benefits in effort-based decision making. Our results add important knowledge toward understanding mechanistic associations underlying complex sedentary behaviors.

## Introduction

In 2007 it was estimated that ~5.3 million global deaths from non-communicable diseases could have been prevented if people engaged in physical activity instead of being physically inactive (Lee et al., 2012). Compounding this further, global statistics show the prevalence of physical inactivity is increasing (Du et al., 2019; Guthold et al., 2018). Over a third of the US population (34.8%) lead sedentary lifestyles (Du et al., 2019; Guthold et al., 2018; Kohl et al., 2012) and the economic burden caused by physical inactivity is estimated to cost private and public health-care systems $53.8 billion per year (Ding et al., 2016, 2017).

To combat the negative consequences of sedentary behaviors, particularly in older adults, the field has studied extensively the beneficial effects of exercise interventions (Gomes-Osman et al., 2018). The most well studied exercise interventions are walking interventions, which are both economical and easily accessible, particularly for older adults. These studies have led to numerous discoveries on the beneficial effects of increased walking on cognitive function, particularly, processing speed, memory and executive function (Erickson et al., 2019). Walking interventions also have been shown to increase hippocampal volume (Erickson et al., 2011) and the plasticity of functional brain networks (Voss et al., 2010). These results are particularly important given these same outcomes are also associated with age-related decline (Buckner, 2004; Luca et al., 2003; Ng et al., 2016; Nobis et al., 2019). However, sedentary behaviors are not simply the inverse of physical activity (Spence et al., 2017; van der Ploeg & Hillsdon, 2017). For example, a person who spends all day sitting at a desk at work but engages in 30 minutes of moderate-to-vigorous physical activity would both be sedentary and engaging in physical activity per World Health Organization guidelines (WHO, 2018). Not all walking interventional studies capture objective measures of sedentary behavior outside of the intervention, yet this information would provide a platform for the study of the determinants of sedentary behaviors within the interventional context. This is particularly important given the determinants of sedentary behaviors are distinct from those of physical activity engagement (Spence et al., 2017). To demonstrate the importance of capturing objective measures of sedentary behavior outside of the intervention, our group has previously shown correlations between objective measures of sedentary behavior and cognition (Burzynska et al., 2020) but more importantly, individual differences in time spent sedentary, regardless of intervention group assignment (Ehlers et al., 2016). Therefore, if one can predict individual differences in future sedentary behaviors early, precision medicine approaches can adapt or prescribe alternative approaches that would reduce the costs of ineffective interventions.

Determinants of sedentary behaviors are likely numerous. Automatic processes, attitudes and habits have been suggested to regulate daily sedentary behavior (Conroy et al., 2013). Association between self-efficacy and sedentary behavior has been shown in meta-analyses (Szczuka et al., 2020) and in younger adults, interventions targeting perceptions of competence and capability (self-efficacy) have been shown to reduce time spent sedentary (Falk et al., 2015). From a cognitive perspective, to successfully overcome short-term costs in favor of longer-term benefits (like reducing sedentary behaviors), executive control functions, such as inhibitory control, flexibility and goal-orientated decision making are required (Cheval et al., 2020).

Extending this further, behavioral choices involving the assessment of motor costs are ever present in day-to-day life and involve the integration of information about available energy resources to weigh physical and motor costs against expected rewards (Klein-Flügge et al., 2016). A theory of energetic cost minimization postulates that we have an innate attraction towards effort minimization whilst maximizing reward (Cheval et al., 2017, 2018; Klein-Flügge et al., 2016; Prévost et al., 2010). This theory is reflected in evolutionary, developmental and situational scenarios, where for example, humans have developed body shapes and neural circuitry refined for energy optimization (Sockol et al., 2007), and during development, energy efficient movements are consolidated through motor practice (Ivanenko et al., 2007), which are constantly adapted in real time to minimize energy costs, such as gait refinements during walking (Selinger et al., 2015). Neural circuitry underlying the valuation of potential behaviors related to physical effort costs have consistently implicated both the anterior cingulate cortex (ACC) and the anterior insula (AI) in these behaviors (Klein-Flügge et al., 2016; Porter et al., 2020; Prévost et al., 2010). For example, in rodents, local field potentials in and coherence between the ACC and the AI correlate with relative performance on a physical effort-based task (Porter et al., 2020). In humans, neuroimaging studies have demonstrated that the ACC is a critical region for decision-making of choices involving motor-costs (Klein-Flügge et al., 2016) and further, that activity in the ACC and the AI represent the devaluation of rewards associated with physical effort (Prévost et al., 2010). Of note, the regions of the ‘ACC’ in these studies would be classified as the mid cingulate (MCC) per sub-region classifications by Vogt (Vogt, 2009).

Discovery of neural predictors of future sedentary behaviors may provide both strong predictive strength as well as mechanistic information relevant for intervention development. The utility and efficacy of functional connectivity to predict future behavioral outcomes has been demonstrated in previous research. For example, Saghayi and colleagues predicted adherence to mental training programs using FC (Saghayi et al., 2020) and Whitfield-Gabrieli and colleagues predicted treatment response in social anxiety disorder with FC, better than clinical measures alone (Whitfield-Gabrieli et al., 2016).

The aims of this present study therefore were to evaluate if baseline measures of FC, executive functions and time spent sedentary could predict future change in objectively measured sedentary behavior, over the course of a 6-month walking intervention in healthy older adults. We hypothesized that FC of regions implicated in both effort-based decision making and inhibitory control (ACC, AI) would be predictive of changes in time spent sedentary, above and beyond behavioral measures.

## Methods

### Participants and study design

Participants in this study participated in a 6-month randomized controlled exercise trial (clinical study identifier: NCT01472744). The study procedures were approved by the University of Illinois Institutional Review Board and written informed consent was obtained from all participants prior to any research activities. Healthy but low active older adults were recruited in Champaign County. Two hundred and forty-seven (169 women) met inclusion criteria for the initial clinical trial, agreed to enroll in the study, and underwent a series of demographic, health, physical activity, neuroimaging and cognitive tests at baseline. Participants in the initial trail were randomized to one of four intervention groups. Of these 247, 72 of these participants were randomized to either the walking intervention group or the walking plus a dietary supplement intervention group (the walking intervention was identical in both groups). For the purpose of this analysis, we chose to analyze these subjects. After excluding six participants for not adhering to more than 50% of the intervention and two participants for high motion artefact in the MRI scan (see below for criteria), 64 participants were ultimately included in this study. We chose to analyze only those in these two intervention groups for two reasons; 1) walking is the most well studied and easily implementable exercise intervention in the literature and 2) a previous analysis with these data showed that there existed intervention-mediated differences in physical activity/inactivity patterns across groups throughout the 6 month (Ehlers et al., 2016), meaning that participants may have had different reasons/mechanisms behind changes in time spent sedentary. For more details on this clinical trial, its primary outcomes and neuroimaging data, please refer to earlier work (Baniqued et al., 2018; Burzynska et al., 2020; Ehlers et al., 2016; Voss et al., 2019). Our current analysis asks a novel question of this dataset that has not been previously assessed. Initially, to enroll in the study, participants must have met the following criteria: were between the ages of 60 and 80 years old, free from psychiatric and neurological illness and had no history of stroke, transient ischemic attack, or head trauma, scored < 23 on the Mini-Mental State Exam, < 21 on a Telephone Interview of Cognitive Status questionnaire and < 10 on the Geriatric Depression Scale, at least 75% right-handed based on the Edinburgh Handedness Questionnaire (a criterion related to functional magnetic resonance imaging (MRI) analyses), demonstrated normal or corrected-to-normal vision of at least 20/40 and no color blindness, screened for safe participation in an MRI environment (e.g., no metallic implants that could interfere with the magnetic field or cause injury and no claustrophobia) and reported to have participated in no more than two bouts of moderate exercise per week within the past 6 months (with the goal of recruiting low active older adults). Table 1 contains complete characterization of the study participants.

### Accelerometry

Time spent sedentary was measured using an ActiGraph accelerometer device (Model GT1M or GT3X; ActiGraph, Pensacola, FL) for one week at baseline and one week post-intervention. Participants were instructed to wear the accelerometer on the nondominant hip during waking hours for seven consecutive days. A valid measurement day consisted of at least 10 hours of valid wear time, with a valid hour defined as no more than 60 consecutive minutes of zero counts with 1-min sampling epochs. Downloaded data (activity counts), represented raw accelerations summed over a given epoch length (60s), which varied based on frequency and intensity of the recorded acceleration (Fanning et al., 2017). Data were processed using cut points designed specifically for older adults (Copeland & Esliger, 2009) such that 50 or fewer counts per minute corresponded with sedentary behavior. (Other cut points included 51–1,040 counts per minute as light physical activity, and 1,041 counts or greater as moderate to vigorous physical activity. Our outcome measure (change in time spent sedentary) was calculated as post-test minus pre-test for the average number of counts defined as sedentary behavior.

### Executive function composite score and Task switching task

As part of the initial clinical trial, participants completed a battery of cognitive tests from The Virginia Cognitive Aging Project (Salthouse & Ferrer-Caja, 2003). A detailed description of each task can be found in a previous open access publication (Baniqued et al., 2018). Of particular interest to the current study, a task-switching test was performed. Successful task switching performance requires a number of executive control functions, such as working memory (holding rule sets in memory), flexibility (respond flexibly to rule changes) and inhibitory control (inhibition of previously appropriate operations and responses (Allport et al., 1999; Rogers & Monsell, 1995)). The task consisted of a number of trials where participants were shown a number between 1 and 9 (except 5) against a colored background. On a pink background, participants were instructed to determine whether the number was odd or even and on a blue background, they were to determine if the number was higher or lower than 5. To best capture flexibility and inhibitory processes, we analyzed performance on the mixed task block (64 practice trials followed by 160 trials) and extracted average accuracy (% of correct responses) and average reaction time from the mixed switch blocks. That is, the accuracy and reaction time from just the blocks where the trails were switched. In addition, to gain a more global measure of executive function we created a composite score (sum of the standardized z-scores) from six executive function tasks (that group together through principal component analysis at the larger study level (Baniqued et al., 2018), which measure multiple forms of abstract, inductive and visuo-spatial reasoning (Shipley abstraction, form boards, letter sets, matrix reasoning, paper folding and spatial relations). ~30% of participants were missing tasks switching data and 1 participant was missing data for the executive function composite score. Accordingly, we implemented data imputation methods using the “mice” package in R using multiple imputed chained equations and the predictive mean matching method (Buuren & Groothuis-Oudshoorn, 2011) (supplementary material section 1), where imputed data replaced missing data for these subjects.

### Magnetic resonance imaging: preprocessing

Participants underwent an MRI scanning session in a 3 Tesla Siemens TIM Trio system with a 12-channel head coil. High-resolution structural MRI scans were acquired using 3D MPRAGE T1-wighted sequences (TR = 1900 ms; TE = 2.32 ms; TI: 900 ms; flip angle = 9°; matrix = 256 × 256; FOV = 230 mm; 192 slices; resolution = 0.9 × 0.9 × 0.9 mm; GRAPPA acceleration factor 2). One run of T2*-weighted resting state echoplanar imaging (EPI) data was obtained with the following parameters: (6min, TR=2s, TE=25ms, flipangle=80°, 3.4 × 3.4 mm^2^ in-plane resolution, 35 4 mm-thick slices acquired in ascending order, Grappa acceleration factor = 2, 64 × 64 matrix).

Preprocessing of the functional resting state data was performed using the CONN-toolbox v.19c (Whitfield-Gabrieli & Nieto-Castanon, 2012), relying upon SPM v.12 (Wellcome Department of Imaging Neuroscience, UCL, London, UK) in MATLAB R2019a (The MathWorks Inc, Natick, MA, USA). The latest default preprocessing pipeline implemented in Conn was performed which consists of the following steps: functional realignment and unwarping, slice timing correction, outlier identification, segmentation (into grey matter, white matter and cerebrospinal fluid) and normalization into standard Montreal Neurologic Institute (MNI) space resampled to 2mm isotropic voxels for functional data and 1mm for anatomical data, using 4^th^ order spline interpolation. Functional scans were spatially smoothed using a 6mm FWHM Gaussian kernel. During the outlier detection step, acquisitions with framewise displacement above 0.9mm or global BOLD signal changes above 5 standard deviations were flagged as outliers using the Artefact Detection Tools (www.nitrc.org/projects/artifact_detect). Two participants were removed from the final analyses for having >40 scan volumes flagged. This cut off was determined based on preserving at least 5 minutes of scanning time (Van Dijk et al., 2009). Additionally, mean motion (framewise displacement) was used as a covariate of no interest in all second level analyses. This was done to be over conservative given previous studies have shown high degree of motion-behavior correlations (Siegel et al., 2017), despite the fact that no motion parameter was significantly correlated with sedentary time in our data (p > 0.05). Denoising of the functional data was performed using a principal component analysis-based correction method, CompCor (Behzadi et al., 2007). Linear regression was used to remove the effects of these artifacts on the BOLD time series for each voxel and each subject taking into account noise components from cerebral white matter and cerebrospinal fluid, estimated subject-motion parameters (3 rotation and 3 translation parameters and 6 other parameters representing their first order time derivatives), scrubbing and constant and first-order linear session effects. Temporal band-pass filtering (0.008-0.09Hz) was applied to remove physiological, subject-motion and outlier-related artefacts. Quality assurance plots of the preprocessing steps are illustrated in supplementary material (figures S2 to S5).

### Seed-based correlations

The average time series in two regions of interest (ROI), the ACC and the right anterior insula (AI) were extracted. We defined our seeds using the 100-parcel functional atlas by Schaefer 2018. Because the functional parcels of the ACC and the rAI in this parcellation extend outside of the anatomical boundaries of interest we limited our seed ROIs to just the functional parcel constrained by the anatomical boundaries of the ACC and the rAI set by the Harvard-Oxford anatomical atlas. This was done by binarizing the parcels from each atlas and using ‘fslmaths’ functions (Functional Magnetic Resonance Imaging of the Brain’s Software Library, http://www.fmrib.ox.ac.uk/fsl) to multiply the two parcels together (see **figure 1A** and **2A** for an illustration of the seed ROIs). Then, Pearson’s correlation coefficients were computed between the average time series in each ROI and the time series of all other voxels in the brain and converted to normally distributed z-scores using Fisher transformation prior to performing the second-level general linear model. Individual change in sedentary time was entered as a covariate of interest in the second-level analysis, controlling for nuisance variables, age, gender, baseline sedentary time and mean framewise displacement, in separate general linear models for each ROI. Results in this second level analyses were estimated using a primary voxel threshold of *p*<0.001 and with family-wise error (FWE) correction threshold of *p*<0.05 at the cluster-level.

**Figure 1.**
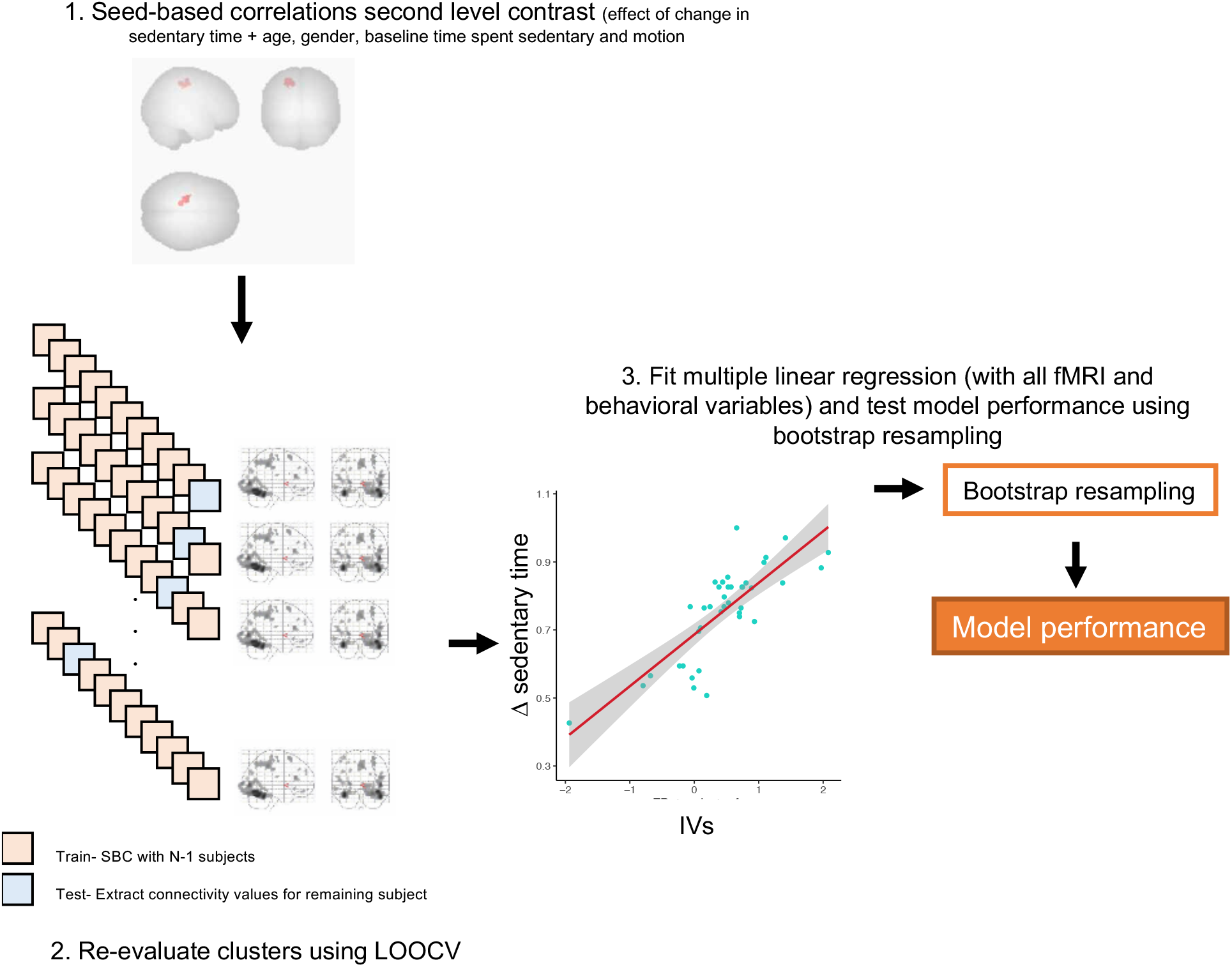
An illustration of the statistical methodology implemented. Seed-based correlations were re-evaluated using LOOCV to provide the prediction performance (R^2^ which represents the squared correlation between the predicted and observed values). Cross-validated FC values were then entered into a multimodal linear regression model with bootstrap resampling to assess the unique contributions of each predictor to the outcome.

**Figure 2.**
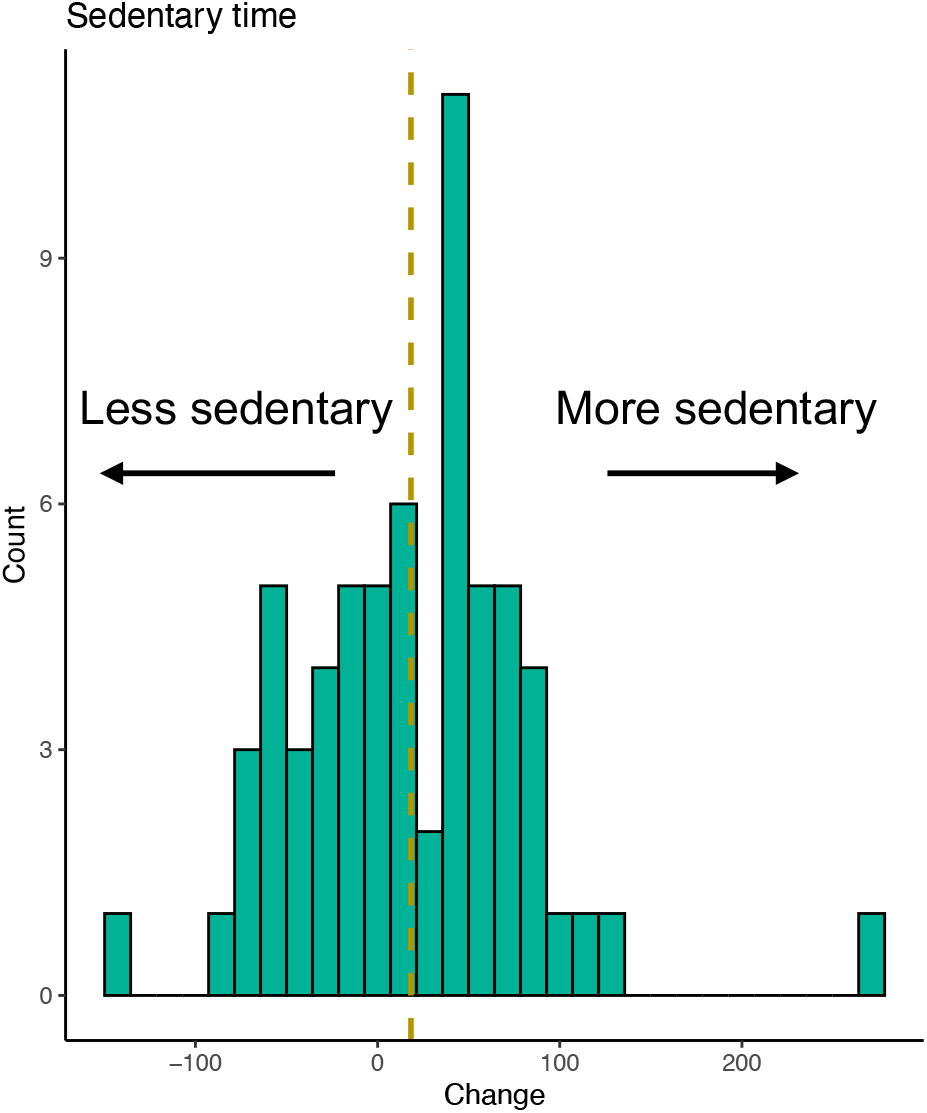
A histogram of participant changes in sedentary time over the 6-month walking intervention.

### Statistical analyses

To assess whether baseline behavioral, cognitive or demographic measures predicted change in time spent sedentary we ran independent linear regression models using leave-one-out cross validation (LOOCV). adjusted R^2^ values are presented which represent the squared correlation between the observed outcome and the predicted values by the model. Model assumptions for linear regression were checked using Q-Q and fitted vs. residual plots in R and the normality of the residuals was formally checked using Shapiro-Wilk tests of normality. The significant influence of outliers was checked using Cooke’s distance with a cut off of 0.5 (no significant outliers removed- see supplementary section 3).

To improve generalizability of the FC results and to minimize potential biases in the voxel selection in our models, significant clusters from functional connectivity seed-to-voxel analyses were re-evaluated using LOOCV. That is, each second level contrast (effect of change in sedentary time, with covariates of no interest (age, gender, baseline sedentary time, mean motion)) was re-ran iteratively leaving one subject out each time. Each iteration then predicted the left-out subjects’ connectivity values with each cluster, resulting in cross-validated connectivity values for each subject. A similar approach was applied in a previous published study (Whitfield-Gabrieli et al., 2016). For each significant cluster we present cross-validated adjusted R^2^ values which represent the squared correlation between the observed outcome and the predicted values by the model.

To assess whether each significant cluster was providing unique explanatory information, we ran a generalized linear model with 500 bootstrapped resamples with cross-validated connectivity values from each of the seed-to-cluster pairs and baseline behavioral and demographic variables. Bootstrap resampling takes random samples from the dataset and provides a distribution of model performance scores (an indication of the variance in the model performance). We chose bootstrapping over other cross validation techniques to reduce overfitting such as k-fold because small sample sizes may result in over-estimation of how the model will generalize given each fold may not be entirely representative of the population being tested. We present the mean adjusted R^2^(which here represents the proportion of variance in the outcome explained by the model predictors) as model performance as well as the distribution of performance across each bootstrap iteration. A depiction of the statistical methodology is outlined in figure 1.

Further, in a complimentary analysis, we dichotomized the participants into either an active (N=25) or sedentary (N=39) group based on whether they decreased Δ<0 or increased Δ>0 their time spent sedentary. We then used logistic regression (with all FC and behavioral/demographic variables) with LOOCV (N-1 participants fit to iteratively predict the out-of-sample participants category) to build cross-validated classifications and estimated accuracy, sensitivity and specificity metrics. LOOCV of SBC clusters was performed in MATLAB using the “spm_crossvalidation” code and all other statistics performed in RStudio Version 3.6.3 (R Foundation for Statistical Computing, Vienna, Austria) using “tidyverse” (Wickham, 2019), “Caret” (Kuhn, 2020) and base R packages.

## Results

Sixty-four low-active healthy older adults (aged 60 to 77 years with a mean of 65 years, 46 females and 18 males) were included in this study. The distribution of the change in time spent sedentary (**Figure 2**) revealed a spread of change where a similar numerical proportion of participants increased time spent sedentary as decreased time spent sedentary.

### Behavioral models

Linear models with leave-one-out cross validation were performed for each baseline behavioral variable and showed that neither baseline measures of time spent sedentary (β = −0.039, SE = .086, *p* = .66, R^2^= −.012), task switching accuracy (β = −5083, SE = 55.19, *p* = 92, R^2^= −.016), reaction time (β = 0.049, SE = .041, *p* = .23, R^2^= .007), nor a composite measure of executive function (β = −1.42, SE = 1.86, *p* = .45, R^2^= −.006), alone were predictive of changes in time spent sedentary.

### Seed-based correlation

The ACC seed revealed a significant positive cluster (voxel *p*<0.001 and FWE cluster-level *p*<0.05 correction) spanning the motor and the supplementary motor areas (peak MNI: −24 −20 +52, size = 425 voxels) associated with change in sedentary time (*T*(58) = 3.47, *p*_fwe_ = <0.001), that is, higher connectivity between the ACC and M1/SMA was associated with an increase in time spent sedentary (**Figure 3C**). Model performance using LOOCV resulted in an R^2^ of .34. Functional connectivity between the rAI and a significant (voxel *p*<0.001 and FWE cluster-level *p*<0.05 correction) cluster in the left temporoparietal/temporooccipital region (areas spanning the middle temporal gyrus, angular gyrus and lateral occipital cortex (peak MNI: −50 −56 +12, size = 194 voxels)) was also associated with change in sedentary time (*T*(58) = 3.47, *p*_fwe_ = <0.001) (**Figure 4C**). Model performance using LOOCV resulted in an R^2^ of .30.

**Figure 3.**
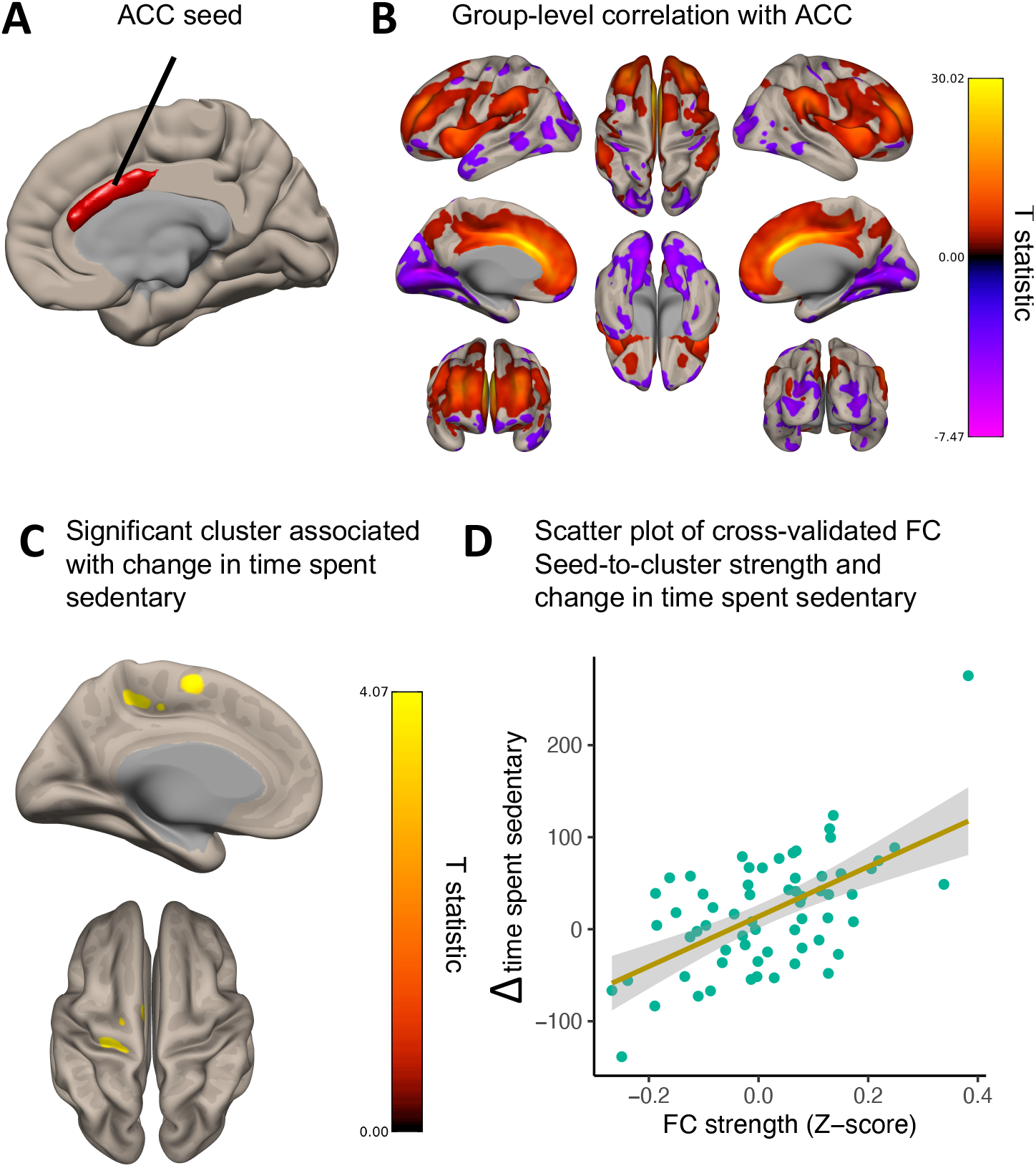
**A.** Illustrates the ACC seed region. **B.** Group-level connectivity with the ACC seed ROI showing functional connectivity with regions of the salience network (bilateral inferior frontal gyrus). **C.** Significant cluster in the left supplementary motor area and the motor area associated with change in time spent sedentary controlling for age, gender, baseline time spent sedentary and mean motion. **D.** Scatter plot of the cross-validated connectivity values (Z) and change in time spent sedentary.

**Figure 4.**
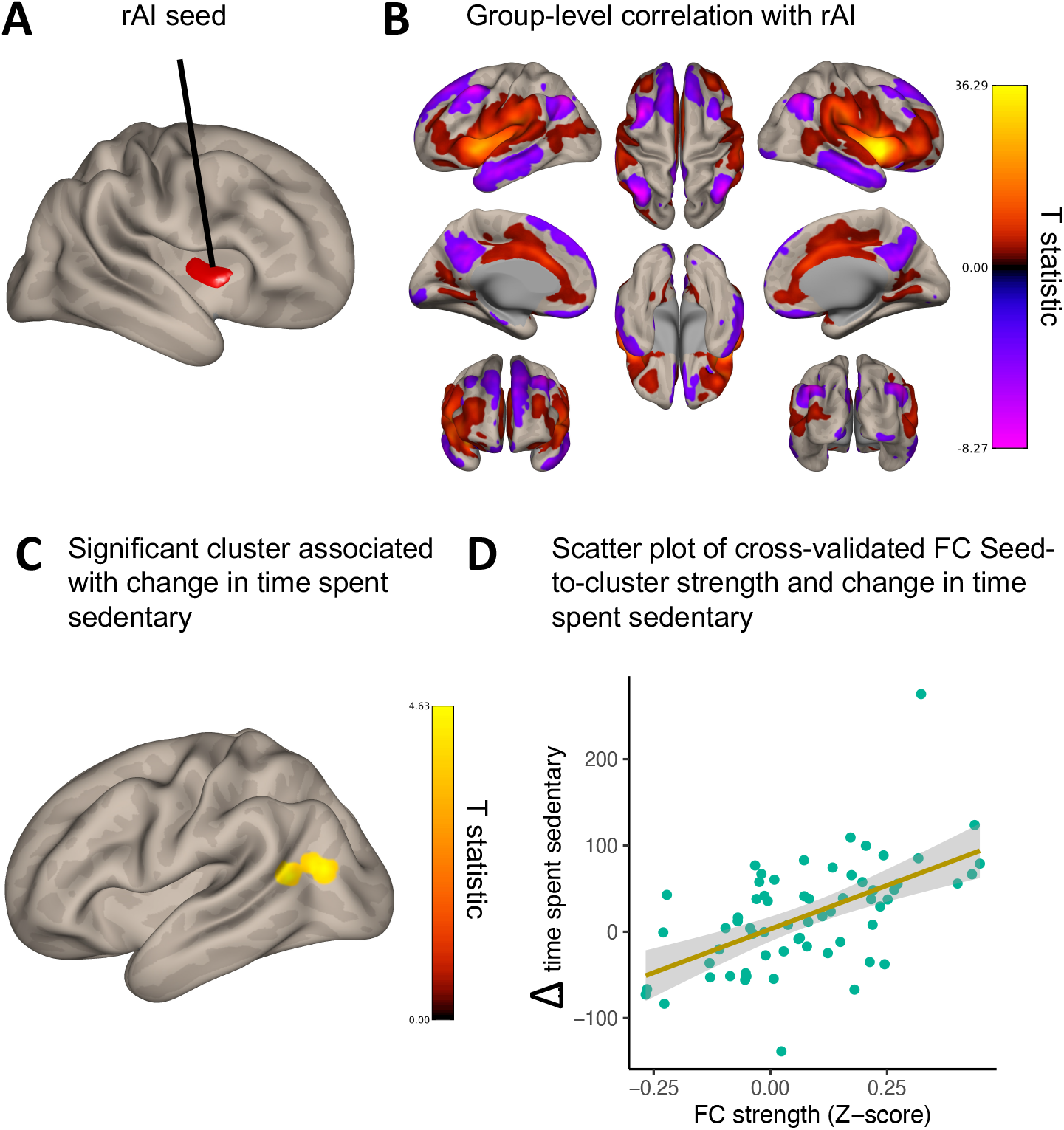
**A.** Illustrates the rAI seed region. **B.** Group-level connectivity with the rAI seed ROI demonstrating our seed functionally connected to the salience network (bilateral insula, temporoparietal junction, inferior frontal operculum, anterior cingulate cortex), and was anticorrelated with the default mode network (inferior parietal lobule, precuneus, superior frontal gyrus). **C.** Significant cluster in the right temporoparietal/temporooccipital region associated with change in time spent sedentary controlling for age, gender, baseline time spent sedentary and mean motion. **D.** Scatter plot of the cross-validated connectivity values (Z) and change in time spent sedentary.

### Multimodal modal

To assess whether each significant cluster was providing unique explanatory information we ran a multiple linear regression model with bootstrap resampling (500 bootstrapped resamples) including cross-validated FC between each seed-cluster pair and baseline variables (baseline time spent sedentary, task switching performance and composite executive function and age and gender). The mean performance of model explained 41% of the variance in the outcome and both FC seed-cluster pairs (ACC: *p* = <.0001, rAI: *p* = <.0001) age (*p* = .0182) and task switching reaction time (*p* = .0139) were significant predictors of change in time spent sedentary (full model results are found in supplementary material). We present a histogram of prediction performance of the model as the variability in the model performance across 500 bootstrap resamples (**Figure 5**).

**Figure 5.**
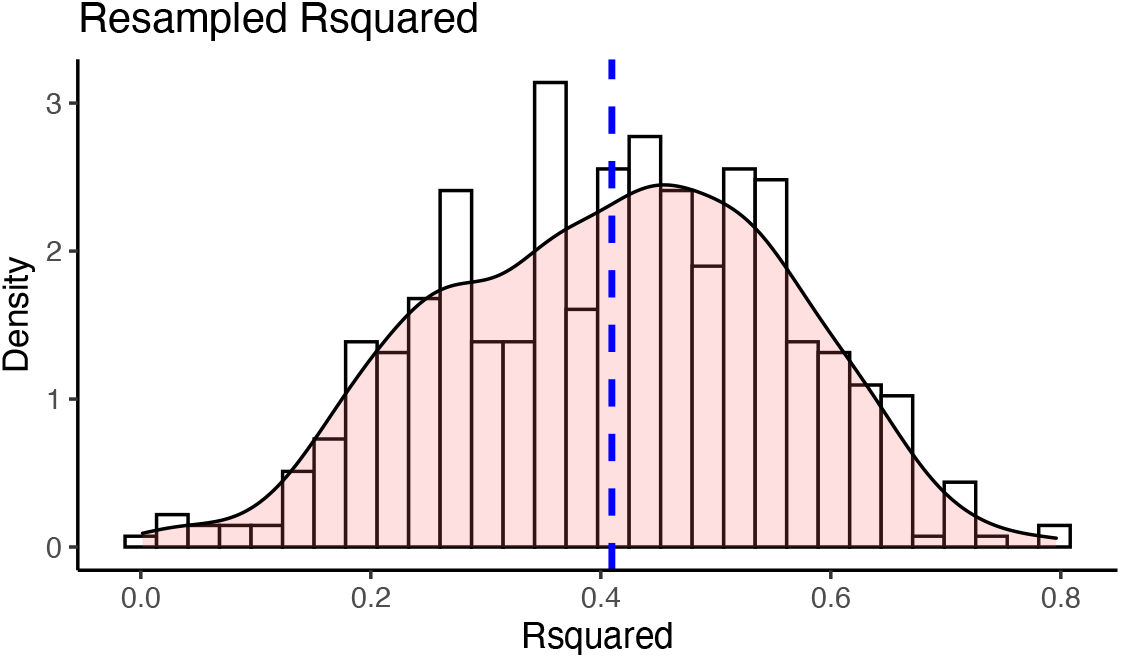
Histogram density plot of the proportion of the explained variance (R^2^) in the outcome from 500 bootstrap resamples of a multiple linear regression model containing cross-validated FC seed-cluster pairs, age, gender, baseline time spent sedentary, executive function and task switching performance as predictors.

### Logistic regression

Logistic regression with LOOCV with FC between each seed-cluster pair and baseline variables (baseline time spent sedentary, task switching performance and composite executive function and age and gender) significantly (95% CI = 0.511,0.757) classified sedentary participants with 64% accuracy, 69% sensitivity and 59% specificity.

## Discussion

The aims of the current study were to assess whether baseline measures of executive function, time spent sedentary and functional connectivity of brain regions implicated in executive control and effort-based decision making could predict changes in time spent sedentary over the course of a 6-month walking intervention. Our results show that while neither baseline measures of executive function nor time spent sedentary could predict changes in time spent sedentary alone, functional connectivity between the ACC and the left SMA/M1 and the rAI and the left temporoparietal/temporooccipital junction were predictive of change in sedentary behavior. In a combined model, baseline reaction time on mixed switch blocks of the task switching paradigm (capturing aspects of inhibitory control and flexibility) was significantly associated with changes in time spent sedentary. Overall, we demonstrate the utility of FC to predict future sedentary behavior and gain mechanistic insight into the executive function mechanisms underpinning this behavior. The following paragraphs discuss our results in light of previous research on FC in the regions shown to be predictive of sedentary time, their relationship with functional networks and the behaviors associated with such networks.

We chose our seed regions (ACC and the rAI) as they have been consistently implicated in effort-based decision making and the integration of motor costs with reward outcomes (Bernacer et al., 2019; Klein-Flügge et al., 2016; Porter et al., 2020; Prévost et al., 2010). Further, these same regions have been implicated in inhibitory control, which has been shown to be important to overcome the posited innate attraction towards effort minimization (Cheval et al., 2020). The function of the ACC and its behavioral role has been highly debated (i.e. does it motivate effortful behaviors? (Mueller et al., 2007; Mulert et al., 2005) or engage in decision-making and deployment of cognitive control? (Kerns et al., 2004)). In an attempt to unify these theories, Holyrood and Yeung (2012) proposed that the ACC supports the selection and maintenance of options and context-specific sequences of behavior directed towards particular goals. In line with this, it has been suggested that poorer monitoring of behavior by the ACC (reflected as increased activity in the ACC during error-related activity in a Go/NoGo task (Hester et al., 2004)) may increase the effort required to inhibit behaviors (Garavan et al., 2006). Highly relevant to our results, one previous study demonstrated that a network involving the ACC and the SMA is critically involved in effort-based decision-making and the integration of motor costs into reward evaluation (Klein-Flügge et al., 2016). More importantly, the same study found that activity in the SMA was stronger in participants who tried to more activity avoid higher efforts (Klein-Flügge et al., 2016). It is plausible therefore that those participants in our study who increased their time spent sedentary were engaging in effort avoidance and/or poor behavioral monitoring, which is reflected as an increase in FC between the ACC (involved in decision making where motor costs are evaluated) and the SMA (has higher activity during effort avoidance). Our ACC seed result also extended into the primary motor cortex (M1) as well (Figure 3C). While voluntary movements and internally-selected actions are more traditionally associated with the SMA (Eccles, 1982) and ACC-to-SMA FC (Mueller et al., 2007), neural projections between the ACC and M1 are present in primates (Morecraft & Hoesen, 1992; Paus, 2001) and in fMRI studies, co-activation of the ACC and motor regions have been seen in working memory tasks (Lenartowicz & McIntosh, 2005). Activity in M1 has been found during mental effort and is likely involved in an attentional network linking behavioral responses to salient stimuli (Otto et al., 2018). Indeed, the left medial portions of the cluster mapped onto the ventral attention network (VAN), a network involved in both attention (Corbetta et al., 2008) and external awareness (Webb et al., 2016). Further, activity in the left motor cortex has been shown to increase as the subjective value of effortful rewards increases (Prévost et al., 2010).

Higher FC between the rAI and a cluster overlapping the left temporoparietal and temporooccipital regions junction (regions covering the superior middle temporal gyrus and the inferior angular gyrus and lateral occipital gyrus) was also predictive of increases in time spent sedentary. The rAI has been proposed to provide an early cognitive control response (Ham et al., 2013) and when mapping this result to a large functional network parcellation (Yeo et al., 2011), both the rAI and portions of this cluster (those in the temporoparietal junction (TPJ)) map onto a broad, bilateral VAN/salience network. Indeed, group level connectivity of the rAI ROI (figure 3B) shows positive FC with salience/VAN regions and is anticorrelated with the default mod network (a hallmark sign of the VAN). The VAN is said to be involved in re-direction of attention to behaviorally relevant stimuli (Corbetta et al., 2008; Corbetta & Shulman, 2002) and is implicated in more external awareness than the synonymous salience network (Webb et al., 2016). Previous research using FC have shown the VAN to be predominantly (but not exclusively) lateralized to the right hemisphere (Fox et al., 2006), nevertheless, bilateral TPJ was confirmed to be part of a broad VAN in a very large (N=1000) study (Yeo et al., 2011). Additionally, the left TPJ’s inclusion in such a network seems to provide a distinct role beyond orientating attention to salient stimuli. For example, Webb and colleagues (Webb et al., 2016) suggested that the left TPJ had a critical role in visual external awareness. The authors suggest that awareness can be disassociated from attention, and that significantly more attention may be drawn to a stimulus when subjects are aware of it (Webb et al., 2016). Another study (Kucyi et al., 2012) suggested that the left TPJ is functionally connected to other regions more associated with executive control and therefore may be more involved in the integration of contextual knowledge about salient stimuli. In accordance, the AI has been suggested to be involved in awareness (Craig, 2011). Further, the AI and the ACC are functionally connected at rest (Medford & Critchley, 2010; Taylor et al., 2009) and across multiple tasks, the AI and the ACC are almost always coactivated (Craig, 2009). Relevant to this study, the broad VAN network of brain regions that are implicated in our seed-based correlations have also been shown to change with advancing age (Deslauriers et al., 2017). Therefore, our results suggest that individual differences in the FC of this broad bilateral VAN, possibly engaging in external awareness, effort-based decision making and effort avoidance, in aging is predictive of changes in time spent sedentary.

Our results can only be interpreted in light of their limitations. The studies that have implicated the brain regions discussed have largely used task-based fMRI whereas we have relied upon intrinsic resting state FC. A future study to prospectively test the role of these brain regions in sedentary behaviors would provide stronger evidence of their mechanistic role. Additionally, participants included in this study were initially recruited as part of a randomized control trial of exercise for cognitive and brain health, whose inclusion criteria required them to be low active (spend < 3 days per week performing physical activity), creating a selection bias for this particular analysis. While this is a limitation, it is likely reflective of the national US population where over a 3^rd^(34.8%) lead sedentary lifestyles (Du et al., 2019). The sample size in our study is relatively small and given the difficulty in objectively measuring sedentary behavior and the cost of running randomized control trials of exercise, we do not have an independent dataset on which to examine the generalizability of these results.

Here we show that individual differences in the baseline FC of multiple brain regions previously implicated in effort avoidance and effort-based decision making predict future change in sedentary time. By understanding the mechanistic correlates of future sedentary behaviors, one can intervene early before spending time and money on ineffective and oftentimes costly interventions. In parallel, steps towards individualized medicine approaches can leverage information about future predictions of behavior change.

## Acknowledgments

We would like to thank Michelle Voss, Anya Knecht, Susan Houseworth, Nancy Dodge, Hilly Tracy, Robert Weisshappel and all of the Lifelong Brain and Cognition and Exercise Psychology Laboratory graduate students and staff for their help in participant recruitment and data collection.

## Ethics approval

The University of Illinois Institutional Review Board approved all procedures used in the study.

## Consent to participate

All participants gave written informed consent before participation in any study procedures, all of which conformed to the Declaration of Helsinki for research involving human subjects.

## Consent for publication

All authors agree to the contents of this manuscript and give consent for its publication.

## Availability of data and materials

All data will be provided upon reasonable request to the corresponding author, without reservation.

## Code availability

Code used in this manuscript includes R syntax for statistical analyses and will be shared upon request to the corresponding author and will eventually be placed on GitHub.

