## Supplementary material for "Resting State Functional Connectivity Predicts Future Changes in Sedentary Behavior"

1. Data imputation

We imputed missing data using the “mice” package in R using multiple imputed chained equations and the predictive mean matching method (Buuren & Groothuis-Oudshoorn, 2011). We ran the impuation 5 times where each creates a different dataset where estimated values replace missing values. 34% of partiicpants were missing both task swithcing measures and one partiicpant the executive function composite measure (supplementary figure 1). Supplementary figure 1 top righ shows the distirburion of imputed data points (blue) overlaid ont the actual data points (red) illustrating the plasubility of the imputed data. Because we are unware of any options to run bootstrap resmapling with pooled data we chose to replace missing data with the imputed data points from the imputed iteration (red density plot lines) with a distribution most similar to the actual dataset (blue density plot line- right) with real values. For task switching accuracy we chose dataset number two (see supplementary figure 1 bottom left) and for task switching reaction time and the one participant for executive function we chose dataset number four (bottom right).

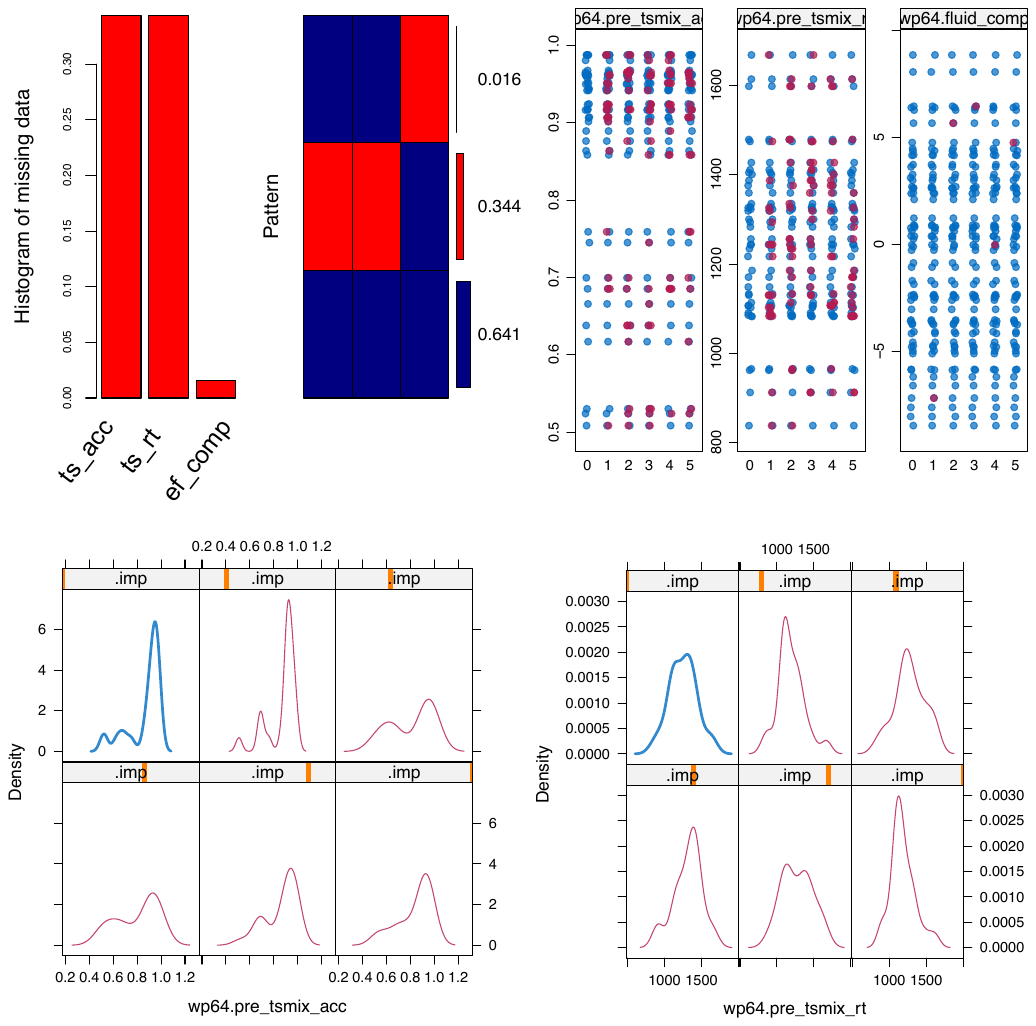

**Supplementary figure 1.** Data imputation results. Imputed dataset numbers in the bottom plots go left to right top to bottom 1-5 with the blue line representing the distribution of the real data points from those without missing data.

2. MRI quality control

The following plots illustrate quality assurance measures of the resting state fMRI data included in this analysis for 64 subjects.

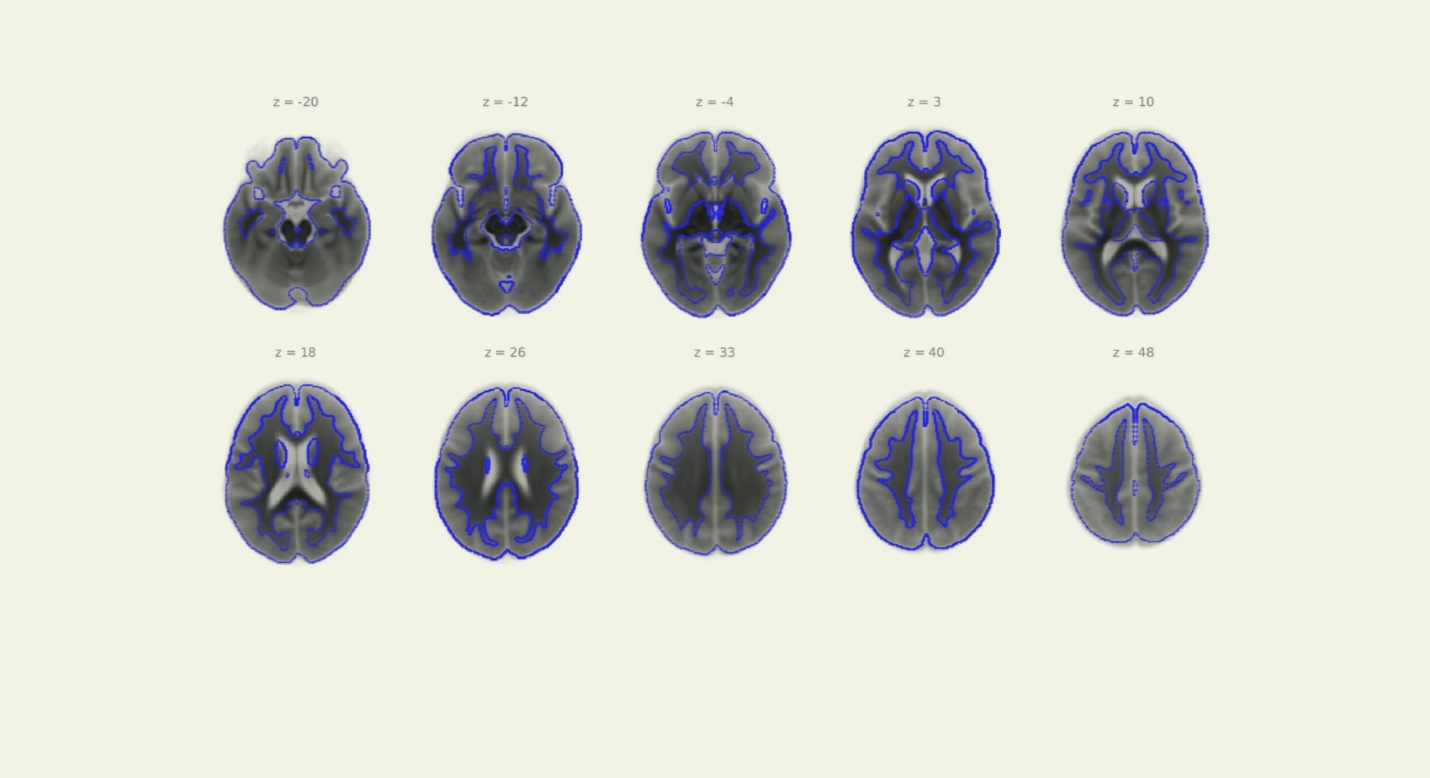

**Supplementary figure 2.** Quality assurance plot, structural normalization.

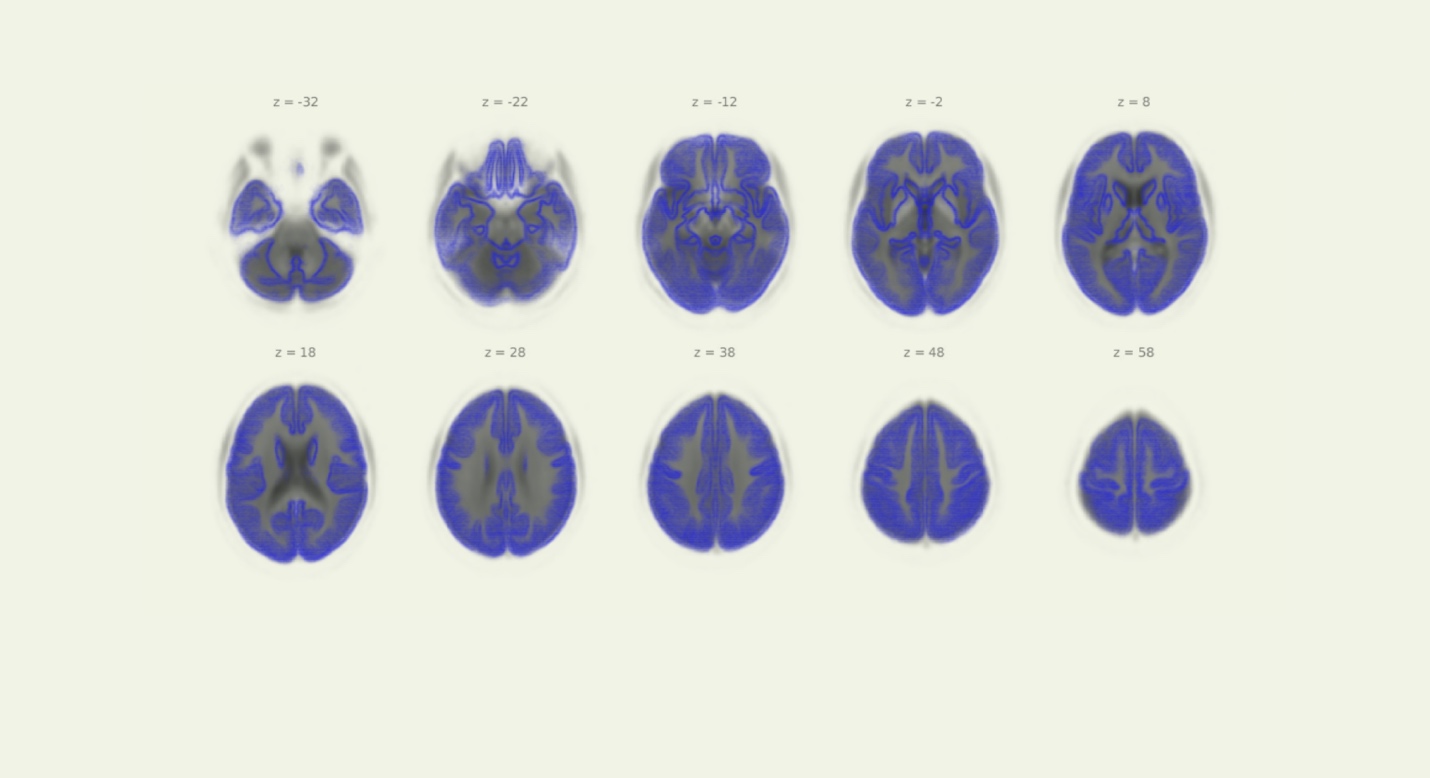

**Supplementary figure 3.** Quality assurance plot, functional data with structural overlay

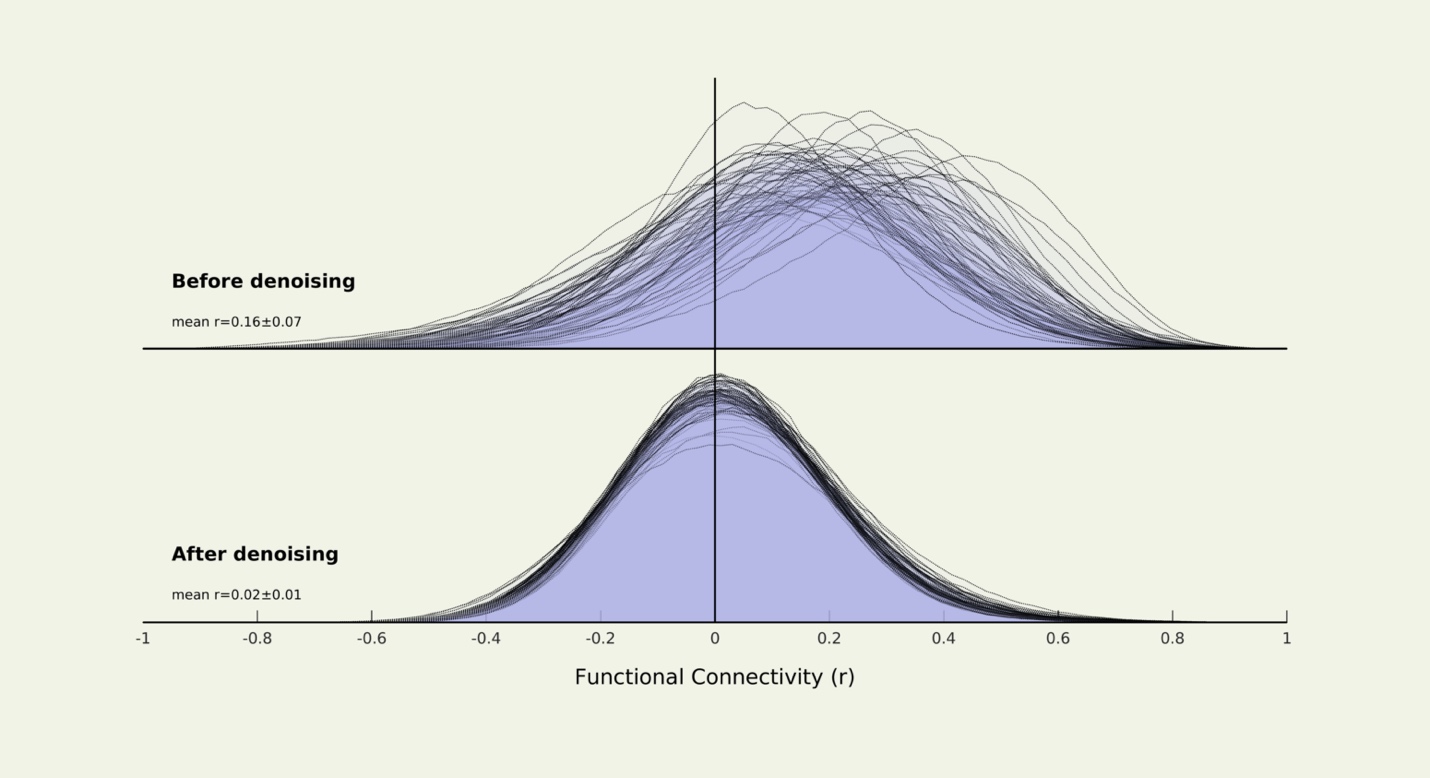

**Supplementary figure 4.** Quality assurance plot, distribution of functional connectivity data before and after denoising

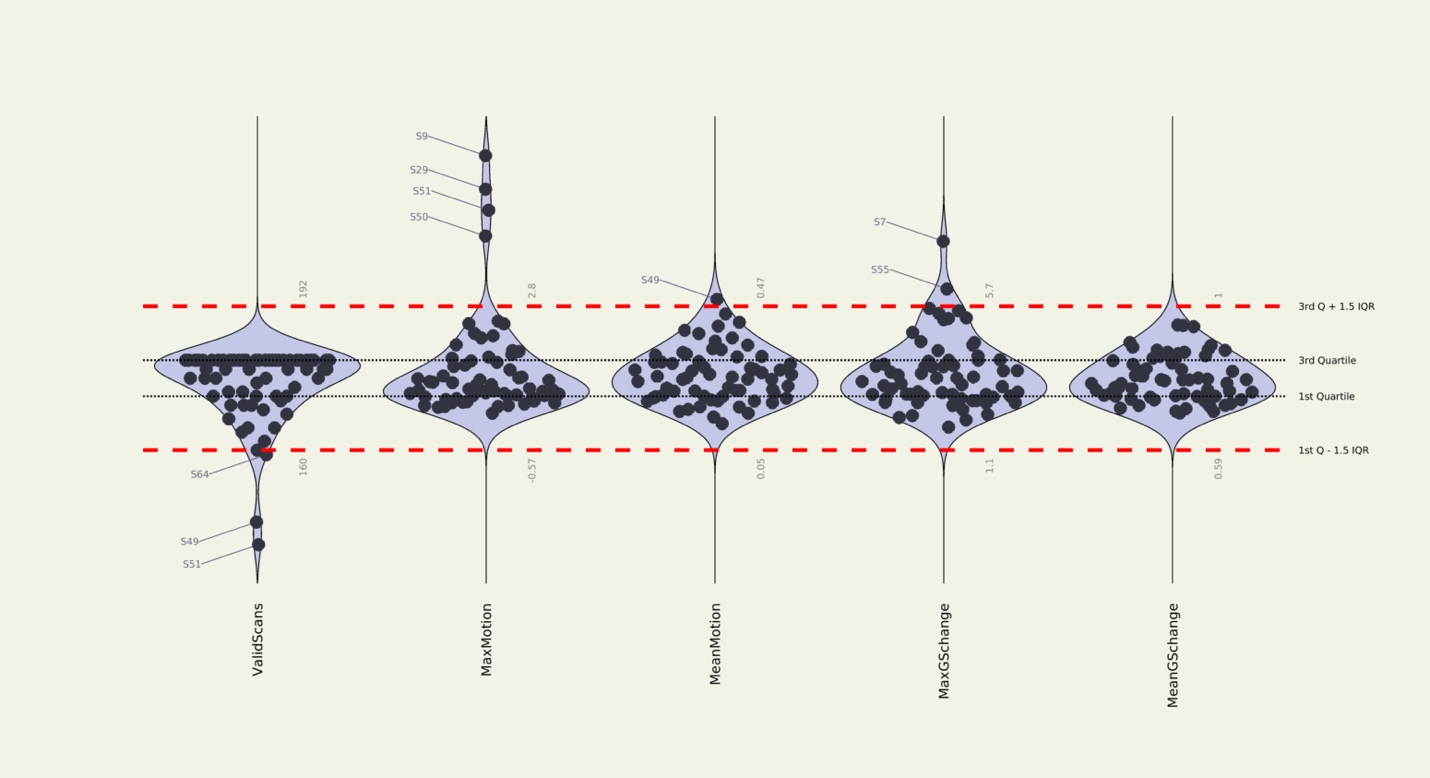

**Supplementary figure 5.** Quality assurance plot, movement parameters

3. Linear model assumptions

Multicollinearity was assessed using the variance inflation factor. No variable in the multiple linear regression with bootstrap resampling had a VIF greater than 1.6 (range = 1.29 – 1.597). A formal test of the normality of the residuals demonstrated their normality (Shapiro wilk p = 0.9). All other model assumptions were checked visually using Q-Q and fitted-vs-residual plots and Cookes distance (cut off of 0.5) for influence of significant outliers (supplementary figure 6).

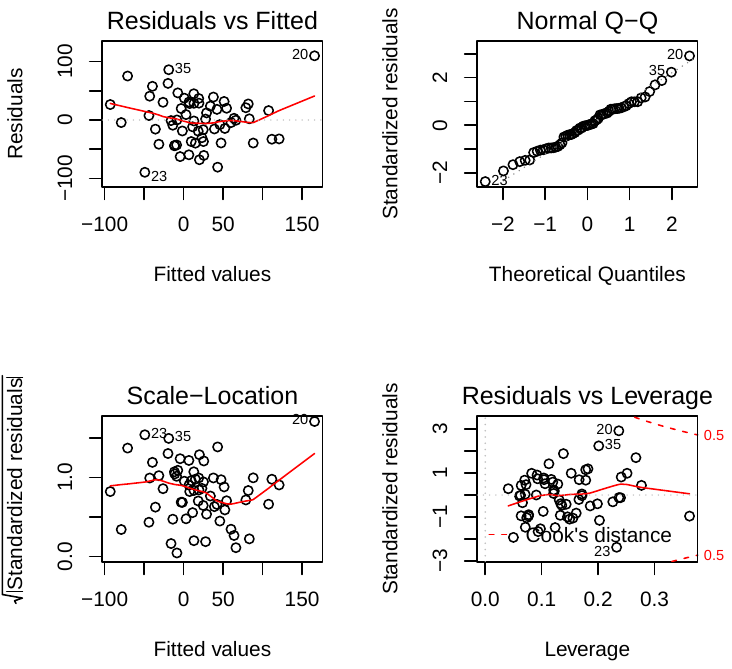

**Supplementary figure 6.** Linear model assumptions.

4. Full bootstrap model coefficients

**Variable	beta	SE	t-value	P-value**

cv_rAI	125.7889	35.8125	3.51	0.0009 ***

cv_acc	278.1258	48.0013	5.79	3.5e-07 ***

age	3.4147	1.4027	2.43	0.0182 *

gender	-23.5951	13.8798	-1.70	0.0948 .

avSedTm0	-0.0470	0.0701	-0.67	0.5048

fluid_comp_imp	-0.6237	1.4817	-0.42	0.6755

pre_tsmix_acc_imp	36.1872	51.1482	0.71	0.4822

pre_tsmix_rt_imp	0.0907	0.0357	2.54	0.0139 *
